# High-throughput genotyping of a full voltage-gated sodium channel gene via genomic DNA using target capture sequencing and analytical pipeline MoNaS to discover novel insecticide resistance mutations

**DOI:** 10.1101/564609

**Authors:** Kentaro Itokawa, Koji Yatsu, Tsuyoshi Sekizuka, Yoshihide Maekawa, Osamu Komagata, Masaaki Sugiura, Tomonori Sasaki, Takashi Tomita, Makoto Kuroda, Kyoko Sawabe, Shinji Kasai

## Abstract

In insects, voltage-gated sodium channel (VGSC) is the primary target site of pyrethroid insecticides. Various amino acid substitutions in the VGSC protein, which are selected under insecticide pressure, are known to confer insecticide resistance. In the genome, the *VGSC* gene consists of more than 30 exons sparsely distributed across a large genomic region, which often exceeds 100 kbp. Due to this complex genomic structure of *VGSC* gene, it is often challenging to genotype full coding nucleotide sequences (CDSs) of *VGSC* from individual genomic DNA (gDNA). In this study, we designed biotinylated oligonucleotide probes from annotated CDSs of *VGSC* of Asian tiger mosquito, *Aedes albopictus*. The probe set effectively concentrated (>80,000-fold) all targeted regions of gene *VGSC* from pooled barcoded Illumina libraries each constructed from individual *A. albopictus* gDNAs. The probe set also captured all homologous *VGSC* CDSs except some tiny exons from the gDNA of other Culicinae mosquitos, *A. aegypti* and *Culex pipiens* complex, with comparable efficiency as a result of the high nucleotide-level conservation of *VGSC*. To enhance efficiency of the downstream bioinformatic process, we developed an automated pipeline to genotype *VGSC* after capture sequencing—MoNaS (<u>Mo</u>squito Na^+^ channel mutation <u>S</u>earch)—which calls amino acid substitutions and compares those to known resistance mutations. The proposed method and our bioinformatic tool should facilitate the discovery of novel amino acid variants conferring insecticide resistance on VGSC and population genetics studies on resistance alleles (with respect to the origin, selection, and migration etc.) in both clinically and agriculturally important insect pests.

## Introduction

Synthetic pyrethroids are currently the most frequently used insecticides for the control of clinically important mosquitos. The mode of pyrethroid’s toxicity is inhibition of the voltage-gated sodium channel (VGSC) in the nervous system (Lund & Narahashi, 1983). Developed resistance against pyrethroids, which is known as knockdown resistance (*kdr*), was first reported in housefly, *Musca domestica*, in 1950s (Busvine, 1951). The *kdr* phenotype as well as another distinct phenotype, super-kdr, was eventually linked to amino acid (aa) substitutions on the two positions, L1014F and L1014F+M918T, respectively, on the gene coding VGSC protein (Miyazaki, Ohyama, Dunlap, & Matsumura, 1996; Williamson, Martinez-Torres, Hick, & Devonshire, 1996). Currently, same aa substitutions as well as various other aa substitutions have been found to be associated with resistance in many medical and agricultural insect pests (Dong et al., 2014; Rinkevich, Du, & Dong, 2013). With this historical background, aa substations in variety of insect species are often projected to corresponding aa position in *M. domestica* VGSC for comparison. Actually, VGSC is highly conserved among insects, and many resistance-conferring aa substitutions are seen parallelly in different species (Davies, Field, Usherwood, & Williamson, 2007). Therefore, it is relatively straightforward to infer the effect of certain aa substitutions in any species if the effect of those substitutions has already been elucidated. However, sequencing analysis of an entire coding sequence (CDS) of the *VGSC* gene from genomic DNA (gDNA) is complicated because *VGSC* genes typically consist of many (>30) small exons sparsely distributed across a large genomic region, which often exceeds 100 kbp Therefore, strategies employing direct sequencing of PCR-amplified genomic fragments usually target only restricted regions where the known resistance-conferring substitutions are frequently found; e.g., IIS5–6 (Dong et al., 2014; Rinkevich et al., 2013). Such a bias may lower the chance of discovering novel resistance mutations existing outside the region investigated.

*Aedes albopictus*, or Asian tiger mosquito, is a medically important mosquito species ubiquitously present on most continents on the Earth. In some regions where the other effective vector, *A. aegypti*, is absent, the species often take a main role for transmitting Chikungunya and Dengue viruses (Kutsuna et al., 2015; Reiter, Fontenille, & Paupy, 2006). The *kdr* substitution in *A. albopictus* had not been reported until the F1534C allele was discovered in Singapore 2009 (S Kasai et al., 2011). Since this discovery, F1534C and other *kdr* substitutions at the same aa position, F1534S and F1534L, were reported from in *A. albopictus* in various geographic locations worldwide (H. Chen et al., 2016; Marcombe, Farajollahi, Healy, Clark, & Fonseca, 2014; Xu et al., 2016). More recently, we also discovered the new *kdr* substitution V1016G in *A. albopictus* by extending the region of the search for mutations (Shinji Kasai et al., 2019).

The next-generation sequencing (NGS) technology has revolutionary reduced the cost and time of DNA sequencing by orders of magnitude. The recent *Anopheles gambiae* 1000 Genomes project, Ag1000G, has uncovered a number of previously unknown nonsynonymous mutations in the *VGSC* gene in *A. gambiae* and *A. coluzzii* (Clarkson et al., 2018), some of which have been suspected to cause resistance directly or indirectly. The study also showed that even neutral variations within or flanking the *VGSC* locus represent valuable information to infer the origin and evolution of resistance. Although whole-genome sequencing may discover novel variants of *VGSC* unequivocally, this naive approach is still too costly per sample just for analyzing *VGSC*. Alternatively, we considered an enrichment approach involving hybridization of oligo DNA/RNA (Gnirke et al., 2009) which is often employed to selectively sequence targeted genomic regions for studies e.g. on genotyping of disease-related genes in humans. This technology is aimed at increasing the depth of reads and the number of samples to be multiplexed per given sequencing capacity in return for limiting the region to be analyzed. In this study, we designed biotinylated oligonucleotide DNA probes from *A. albopictus VGSC* CDSs. The probe set efficiently concentrated targeted regions from the gDNA of individual *A. albopictus*. Although the probe set was designed from the *A. albopictus VGSC* gene, the same probe set captured most CDSs of *A. aegypti* and *Culex pipiens* complex as a result of the high nucleotide conservation of *VGSC*. This technology allows for full-CDS analysis of the complex *VGSC* gene in a relatively low-cost and highly multiplexed manner, which is expected to promote discoveries of novel resistance-conferring aa substitutions both in medical and agricultural insect pests.

## Materials and Methods

### Design of custom probes

The full-length *VGSC* gene (AALF000723-RA in gene set: AaloF1.2) was found in scaffold JXUM01S000562 in the genome assembly of an *A. albopictus* Foshan strain, AaloF1 (X.-G. Chen et al., 2015) hosted on vectorbase.org (Giraldo-Calderón et al., 2015). Because the annotation missed some CDSs (entire exon 19c for instance), we refined the annotation by aligning shotgun-sequenced NGS reads of *VGSC* cDNA (Shinji Kasai et al., 2019) using Hisat2 (Kim, Langmead, & Salzberg, 2015) and the *M. domestica* VGSC protein sequence (GenBank accession No.: AAB47604) via BLASTX (Altschul, Gish, Miller, Myers, & Lipman, 1990). Compared to AaloF1.2, the refined annotation included three added CDSs, and four extended CDSs (see detail in Table S1). Among the 35 coding CDSs in total, sizes of 34 (32 + 2 mutually excluding exons) CDSs matched to the *A. aegypti VGSC* CDSs annotated by Davies et al. (2007). Therefore, the numbering of exons in this paper was set to be concordant with the *A. aegypti VGSC* exons described by Davies et al. (2007). The additional optionally used 45 bp small exon, referred to as exon 16.5 here, was found in cDNA data between exons 16 and 17 in the genome. All CDS sequences, some of which contained a flanking intronic region (for tiny exons less than 120 bp in size) were submitted to the IDT website (https://sg.idtdna.com) to design 120 bp xGen Lockdown biotinylated oligonucleotide DNA probes with2x Tiling density option. We also included some exons of other genes flanking VGSC (AALF020128, AALF020129, AALF020130, AALF000725, AALF000726, AALF000727, AALF000728, and AALF000730) or a gene nested in the intronic region of *VGSC* (AALF020132) during the probe design to take advantage of the population genetic analysis in future studies. From 15 kbp genomic regions in total, 229 probes were designed (Table S2), of which 145 target *VGSC*. Nonetheless, in the more recent contiguous assembly of the C6/36 cell line (see below), AALF020132, AALF000725, AALF000726, AALF000727, AALF000728, and AALF000730 are not located in the same assembly with the *VGSC* locus. For this reason, in this paper, we evaluate the performance of the probe set only in terms of *VGSC* CDS enrichment.

### Samples

Fifty-six mosquitos either belonging to species *A. albopictus, A. aegypti* or *Culex pipiens* complex—either kept in the laboratory or caught in the wild (Table 1)—served as a source of gDNA. Of those, strains Aalb-SP, Aaeg-SP, and Cpip-JPP were already known to possess haplotypes with 1534C, 989P-1016G, and 1014F aa variants, respectively (Hardstone et al., 2007; S Kasai et al., 2014; Shinji Kasai et al., 2019).

**Table 1.** Mosquito samples used in this study

| ID | Species | Lab. Colony<br>/ Wild | num | Description | DDBJ<br>Accession no. |
| --- | --- | --- | --- | --- | --- |
| Aalb-SP | *A. albopictus* | Lab. Colony | 8 | Originated from Singapore in 2016. This strain is known to possess 1534C *kdr* variant (Kasai et al., 2019). | DRR167925–932 |
| Aalb-Viet | *A. albopictus* | Lab. Colony | 2 | Originated from Hanoi, Viet Nam in 2016. | DRR167935–936 |
| Aalb-Okayama | *A. albopictus* | Lab. Colony | 2 | Originated from Okayama, Japan in 2015. | DRR167923–924 |
| Aalb-Toyama | *A. albopictus* | Lab. Colony | 2 | Originated from Tokyo, Japan in 2015. | DRR167933–934 |
| Aalb-Ishigaki | *A. albopictus* | Lab. Colony | 2 | Originated from Ishigaki-zima, Okinawa, Japan in 2016. | DRR167921–922 |
| Aalb-Yona | *A. albopictus* | Wild | 8 | Caught wild from Yonaguni-zima, Okinawa, Japan in 2017. | DRR167937–944 |
| Aaeg-Mex | *A. aegypti* | Lab. Colony | 16 | Originated from Monterrey, Mexico in 2008. | DRR167897–912 |
| Aaeg-SP | *A. aegypti* | Lab. Colony | 8 | Originated from Singapore in 2009. This strain is known to possesses 989P–1016G *kdr* haplotype (Kasai 2014). | DRR167913–920 |
| Cpip-JNA | *C. quinquefasciatus* | Lab. Colony | 2 | This strain was selected for *CYP9M10* genotype (Itokawa et al., 2011) from JHB strain originated from Johannesburg, South Africa in 2001 (Arensburger et al., 2010). | DRR167945–946 |
| Cpip-JPP | *C. quinquefasciatus* | Lab. Colony | 2 | Originated from Saudi Arabia, selected by permethrin for 20 generations (Amin et al., 1989). This strain is known to possesses 1014F *kdr* variant (Hardstone et al., 2007) . | DRR167947–948 |
| Cpip-Ryo | *C. pipiens pallens* | Lab. Colony | 2 | Originated from Kanagawa, Japan in 2015. | DRR167951–952 |
| Cpip-JP_mix | *C. pipiens* form *molestus* | Lab. Colony | 2 | Mixed from several lab. colonies originated from different places of Japan in 2003–2004. | DRR167949–950 |

### gDNA extraction

gDNA was individually extracted from the whole body of an adult or pupa using the MagExtractor Genome Kit (TOYOBO). The protocol was modified to conduct the extraction in 8-strip PCR tubes or a 96-well PCR plate as follows. The whole body of a single insect was homogenized in a PCR well containing 50 μl of the Lysis & Binding Solution and zirconia beads (ø 2 mm; Nikkato) in TissueLyser II (QIAGEN) at 25 Hz for 30 s. After that, the samples were centrifuged at 2000 x *g* for 1 min to precipitate large debris, and each supernatant was transferred to a new well containing 50 μl of the Lysis & Binding Solution and 5 μl of DNA-binding Magnetic Beads. The solution was shaken in MicroMixer E-36 (TAITEC) at the maximum speed (2500 rpm) for 10 min, and then, on a magnetic plate, the supernatant was discarded. The beads bound to DNA were washed twice with 100 μl of the Washing Solution and twice with 75% ethanol each. Finally, DNA was eluted with 50 μl of low-TE buffer (0.1 mM EDTA, 10 mM Tris-HCl pH 8.0) by shaking in MicroMixer E-36 at the maximum speed for 10 min. The obtained DNA was quantified with the Qubit Highly Sensitive DNA Assay Kit (Invitrogen). The obtained DNA concentration ranged from 2.3 to 8.6 ng/μl for *A. albopictus*, 5.3 to 10 ng/μl for *A. aegypti*, and 7.8 to 11 ng/μl for *C. pipiens* complex mosquitos.

### Library construction and hybridization capture

Illumina libraries with TruSeq barcode adapters were prepared using NEBNext Ultra II FS DNA Library Prep Kit for Illumina (NEB) on the 1/4 scale of the manufacturer-suggested protocol. Briefly, 4 μl of the gDNA extracted above (without adjusting the concentration) was mixed with 0.4 μl of the Enzyme Mix, 1.4 μl of Reaction Buffer, and 1.2 μl of H_2_O on ice. The mixture was then incubated at 37 °C for 10 min followed by incubation at 65 °C for 30 min. Those end-prepped DNAs were directly ligated with Illumina adapters by the addition of 0.5 μl of TruSeq 96 dual-index adapters (Illumina) instead of adapters supplied with the kit, 6 μl of the Ligation Master Mix, and 0.2 μl of Ligation Enhancer, with incubation at 20 °C for 15 min. Next, the libraries were incubated at 65 °C for 30 min to inactivate the ligase; then, all the 56 libraries were pooled together in a single 1.5 ml LoBind tube (Eppendrof). The pooled library was purified with 1.2x SPRIselect (Beckman Coulter) and eluted with 20 μl of low-TE buffer. A 7 μl aliquot of the pooled library was aliquoted and mixed with 0.8 μl of a 10 μg/μl UltraPure Salmon Sperm DNA Solution (Invitrogen) in a PCR-tube. Then. the mixsture was concentrated by incubation at 80 °C for 10 min while the lid of the tube and thermal cycler were opened. The concentrated library mix was hybridized, captured, and washed with the designed oligo DNA probe set and the xGen Hybridization and Wash Kit (IDT). After that, the streptavidin magnetic beads were subjected to PCR amplification with HiFi Kapa (Kapa Biosystems) for 12 cycles. The amplified library was purified with 1.2x SPRIselect beads and quantified by real-time PCR using P5 and P7 adapter primers and qPCR double quencher probe (6-FAM)-5′-ACACTCTTT-(ZEN)-CCCTACACGACGCTCTTC-3′-(Iowa Black FQ) (IDT) in the PrimeTime Gene Expression Master Mix (IDT). Serial dilutions of the phiX library (Illumina) were used for construction of the standard curve. The quantified library was sequenced on Illumina MiniSeq with the Mid Output Kit (Illumina) for 151 cycles from both ends along with the libraries from other studies.

### Reference genomes and annotation for *VGSC*

Although the probe sets were designed from assembly AaloF1, we chose a C6/36 cell line genome assembly, canu_80X_arrow2.2 (Miller et al., 2018), as a reference genome of *A. albopictus* for further bioinformatic analysis because this assembly has better contiguity and fewer scaffolds than AaloF1 does. In the canu_80X_arrow2.2 assembly, the whole *VGSC* gene was found in scaffolds MNAF02001058.1 and MNAF02001442.1 annotated as Gene IDs LOC109421922 and LOC109432678, respectively, in the NCBI *Aedes albopictus* Annotation Release 101. The two *VGSC* genes were assumed to be redundant haplotigs. To avoid dual mapping of the NGS reads, we purged MNAF02001442 by hard-masking this entire scaffold (replacing all bases with the “N” character) rather than MNAF02001058.1 because LOC109432678 in MNAF02001442.1 has a single frame-shifting nucleotide deletion in the thymine homopolymer track within exon 4 (TTTTTT → TTTTT), which was suspected due to an uncorrected base-calling error. LOC109421922 was defined by the number of transcriptional variants in the NCBI’s annotation because *VGSC* is known to have complex alternative splicing patterns (Davies et al., 2007). Nevertheless, we simplified the *VGSC* gene model into two possible transcriptional variants to build a GFF3 annotation file for annotating aa changes. These two transcripts include all the regions of mandatory or optional CDSs but differ by the two mutually exclusive exons “19c/k” and “26d/l,” where one carries exons “19c” and “26k,” and the other contains exons “19d” and “26l.” CDSs of all the transcriptional variants of LOC109421922 were merged via overlaps. Those merged CDSs perfectly matched AaloF1 except for LOC109421922 including exon 16.5 and except for one mutually exclusive exon “26k” whose sequence itself was found to be intact in MNAF02001058.1.

The *VGSC* gene in the chromosome level assembly of the *A. aegypti* genome, AaegL5.0 (Matthews et al., 2018), was annotated in the same manner. The whole *VGSC* gene is encoded as AAEL023266 (the NCBI *Aedes aegypti* Annotation Release 101) on chromosome 3. AAEL023266 has 13 transcripts, these CDSs were merged via overlaps as in the canu_80X_arrow2.2 assembly of *A. albopictus*. AAEL023266 appeared to be lacking an exon corresponding to exon 16.5, whereas we found its sequence between exons 16 and 17. AAEL023266, however, contains an additional exon between exons 11 and 12. The 21 bp small exon was assumed to correspond to exon “12” in the *Anopheles gambiae* genome described by Davies et al. (2007) and is situated within the intracellular loop between domains I and II. We found the sequence homologous to this exon also in the two *A. albopictus* genome assemblies, AaloF1 and canu_80X_arrow2.2; thus, which means we had failed to include this exon in the probe design. The complete *VGSC* sequence was also found in scaffold NIGP01000811 and was assumed to be a redundant haplotigs. This scaffold was purged from the assembly by hard-masking.

In *C. quinquefasciatus* genome assembly Cpip_J2 (Arensburger et al., 2010), the *VGSC* gene is located in scaffold supercont3.182. The *VGSC* gene in supercont3.182, however, contains shorter exon 13 which was truncated by scaffolding gap and lacks the entire exon 14. Complete exons 13 and 14 were found in another scaffold, supercont3.1170, which contains an incomplete *VGSC* gene probably as a haplotig. We fused contig AAWU01037504.1 containing exons 13 and 14 of *VGSC* from supercont3.1170 into supercont3.182 to restore the complete coding sequence of the *VGSC* gene, thereby creating supercont3.182_2 (Fig. S1A). The *VGSC* in supercont3.182 (and supercont3.182_2) still contained *kdr* aa substitutions, L932F and I936V, as already reported by Davies et al. (2007) plus unusual frameshifting deletions in exons 26l and 32 (Fig. S1B). For these reasons, supercont3.182_2 was further polished by the *consensus* module in BCFtools (Danecek & McCarthy, 2017) with the “-H 1” option using the variant information for Cpip-JNA-01 in the VCF file generated as described below, thereby finally resulting in supercont3.182_3. The latter scaffold was added to the genome assembly, and the original scaffolds supercont3.182 and supercont3.1170 were purged by hard-masking. We were not able to find an exon corresponding to “exon 16.5” in *A. albopictus* and *A. aegypti*.

### Bioinformatic analysis

The FASTQ data were mapped to the reference using *BWA mem* (v.0.7.17) (Li & Durbin, 2009) with default options. The resultant BAM files were sorted by the *sort* program from the SAMtools suite (v.1.9) (Li et al., 2009), and we removed PCR duplicates by the *rmdup* programs from the SAMtools. Variant calling was performed on the resulting BAM files of each species in the FreeBayes software (v.1.2.0) (Garrison & Marth, 2012) with default options. The variant annotation (for aa changes) was conducted with the *csq* program from the BCFtools suite (v.1.9) (Danecek & McCarthy, 2017) with options “*-l -p a*”. Finally, the discovered aa changes were projected onto the corresponding position in the *M. domestica* VGSC protein sequence (GenBank accession No.: AAB47604). Those bioinformatic processes (Fig. 1A) were automated in pipeline tools *MoNaS* (<u>Mo</u>squito <u>Na</u>^+^ channel mutation <u>S</u>earch; https://github.com/ItokawaK/MoNaS) written in the Python3 script language.

**Fig. 1.**
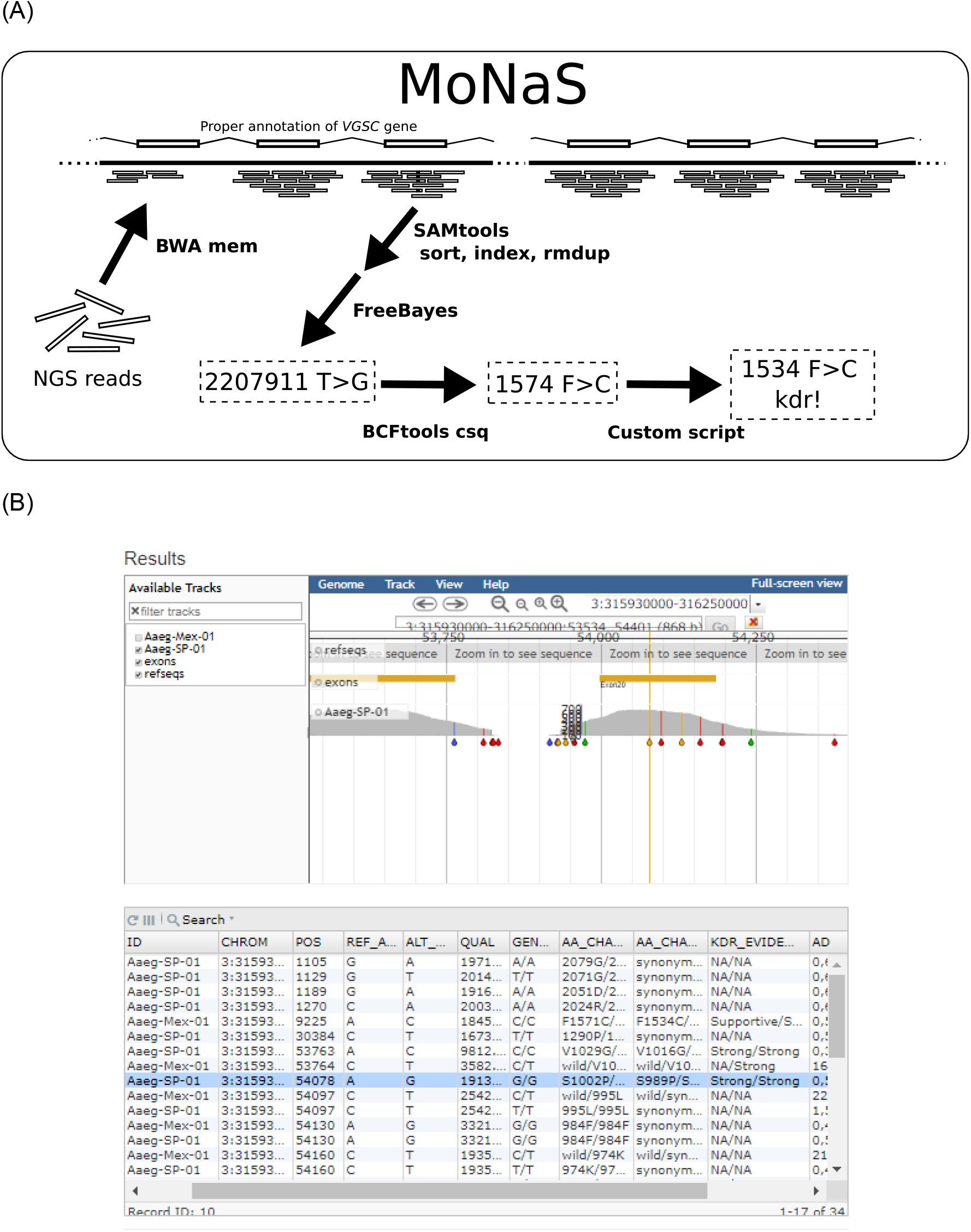
Analytical pipeline MoNaS. (A) A diagram for analytical pipeline MoNaS. MoNaS executes several bioinformatic tools to call variants and aa substitutions. Finally, a custom script converts species aa coordinates to the standard housefly aa coordinates, tells whether each aa substitution is among the known listed *kdr* substitutions and creates a human-readable table from Variant Call Format (VCF). (B) Image of the result output page from MoNaS web-service.

For estimating the level of enrichment, five sets of random data on whole-genome shotgun paired-end reads (150 bp x 2, 300 ± 50 bp insert length, 1 million read pairs) from each reference genomic assembly were simulated in the *wgsim* software (https://github.com/lh3/wgsim). The *multicov* program from the Bedtools suite (v.2.27.1) (Quinlan & Hall, 2010) was applied to calculate the number of reads overlapping with any targeted CDS regions. Nucleotide identities of exons were calculated using *Muscle* (Edgar, 2004) and BioPython’s *Phylo* package (Talevich, Invergo, Cock, & Chapman, 2012). *R* (v.3.5.1) (R_Development_Core_Team, 2014) and the *ggplot2* package (Wickham, 2016) were utilized for summarizing and visualizing the data.

## Results

For the 56 mosquito gDNA samples, 4.9 million demultiplexed read pairs (150 bp PE) in total were obtained after a single run of Illumina MiniSeq. From those samples, 40–170, 50–120, and 63–100 thousand read-pairs were obtained from each individual mosquito of species *A. albopictus* and *A. aegypti* and *C. pipiens* complex, respectively. The raw read data were deposited to DDBJ Sequence Read Archive (DRA) under BioProjectID: PRJDB7889. In *A. albopictus*, 44% of all reads on average overlapped with any of the *VGSC* CDSs under study (<u>R</u>eads overlapping <u>p</u>er <u>k</u>ilobase exon and per <u>m</u>illion sequenced reads: RPKM = 6.5 x 10^4^), which was approximately 8.3 x 10^4^-fold enrichment compared to simulated random data on whole-genome shotgun sequencing (Fig. 2). Although the probe set was designed based on the *A. albopictus* genomic sequence, the same probe set captured *VGSC* CDSs from the gDNA of *A. aegypti* and *C. pipiens* complex at an on-target rate similar to that of *A. albopictus* (Fig. 2).

**Fig. 2.**
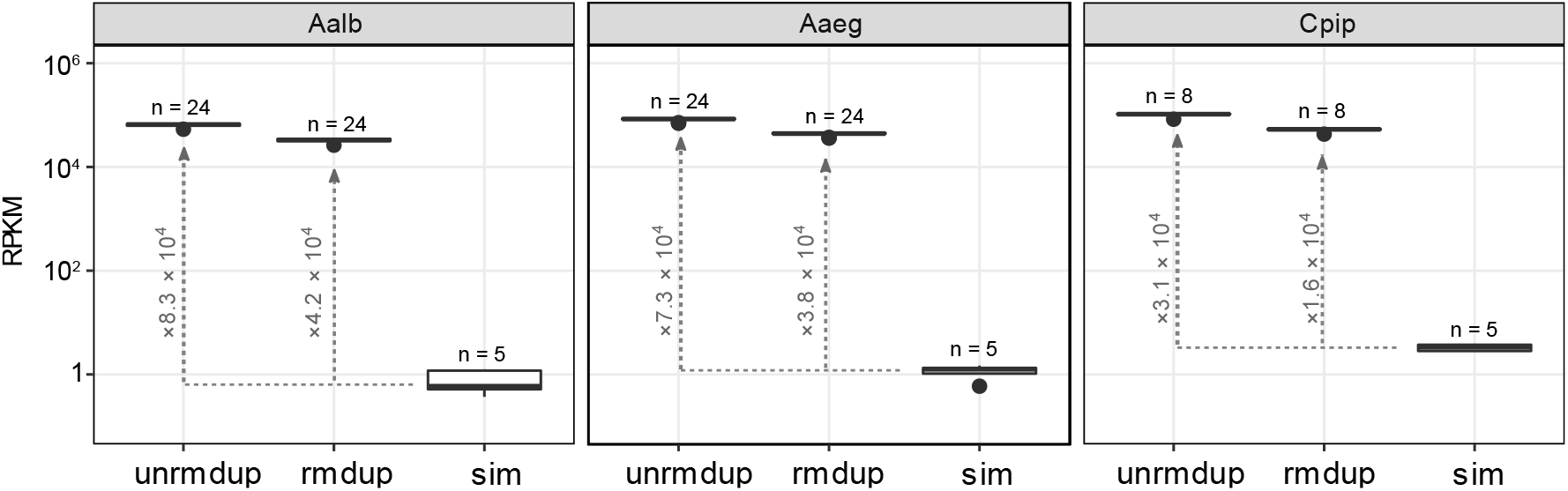
The NGS reads are enriched in targeted *VGSC* exons by capture. Distributions of RPKM (number of sequencing reads overlapping to the targeted *VGSC* exons <u>p</u>er 1 <u>k</u>bp total exon length and one <u>m</u>illion reads) for *A. albopictus* (Aalb), *A. aegypti* (Aaeg) and *C. pipiens* complex (Cpip). Labels “unrmdup” and “rmdup” indicate before and after removal of PCR duplicates, respectively. Label “sim” indicates simulated whole genome shotgun (WGS) reads randomly drawn from genome of each species (replicated five times in each species). The values associated with dotted line with arrowhead indicate sizes of fold-change (levels of enrichment) compared to simulated WGS data.

Fig. 3 shows a distribution of median and minimum sequencing depths within each CDS in each individual sample after PCR duplicates were removed. In *A. albopictus*, most nucleotides in all exons were covered deeply with minimum bias in all samples. In *A. aegypti* and *C. quinquefasciatus*, however, some exons were covered at relatively low depth partly or entirely. In particular, exons 2 and 16.5 were covered at nearly or absolutely zero depth.

**Fig. 3.**
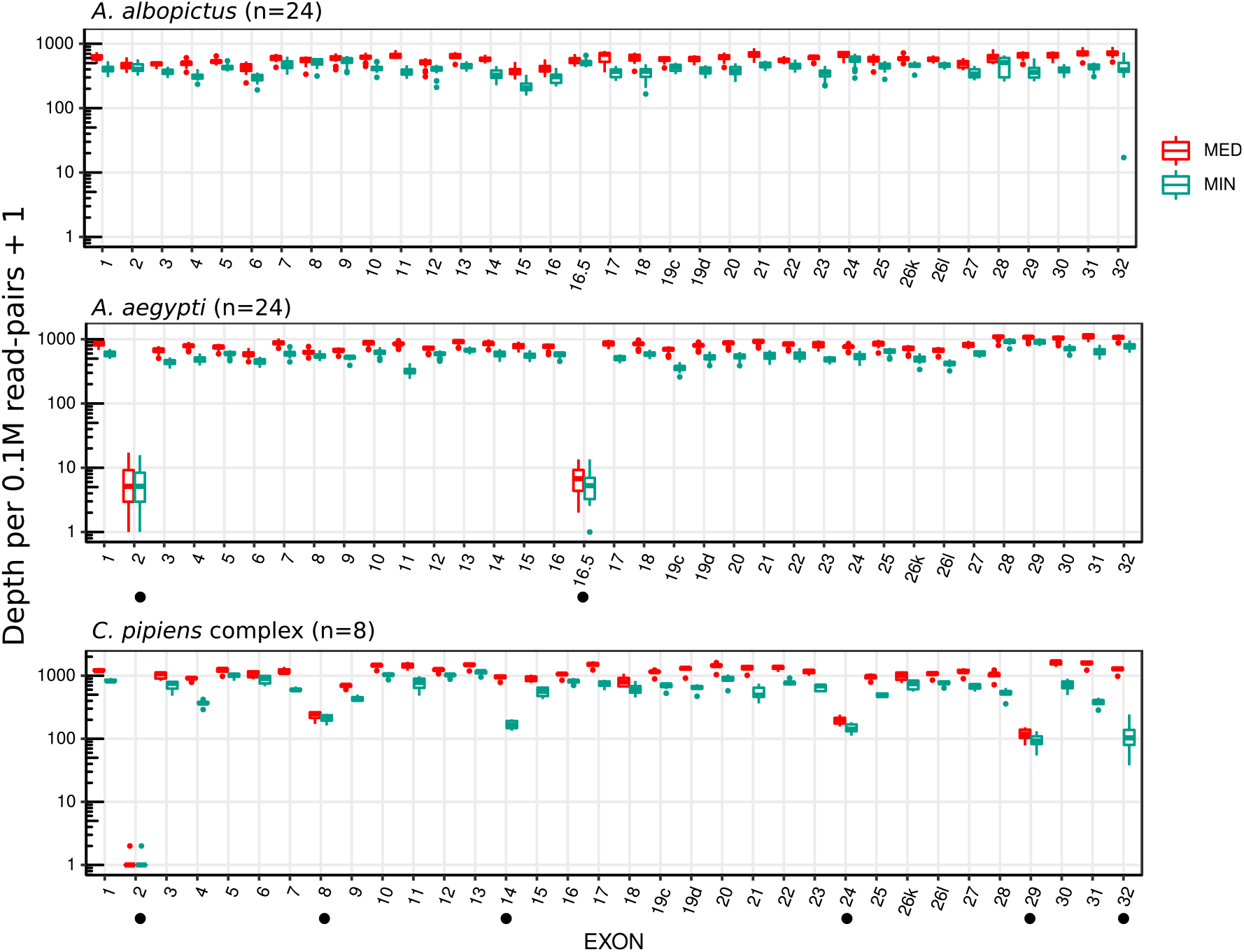
Coverage of targeted *VGSC* exons. Distribution of median (MED) and minimum (MIN) depths per nucleotide (on a logarithmic scale) within each exon and each individual sample after PCR duplicates were removed. Exons labeled with “●” contained nucleotide sites with relatively low coverage.

Fig. 4 presents a distribution of the allele balance in genotypes containing single or multiple nucleotide variants (SNVs or MNVs). The ratio of the first allele in a heterozygous genotype was distributed mostly around 50%, which was substantially different from the homozygous genotype (near 100%) except for one SNV or MNV site in exon 32 of the *A. albopictus* gene located in the GGT (Gly) trinucleotide tandem repeats variable in length near the C-terminus of VGSC, where accurate calling of the genotype is difficult.

**Fig. 4.**
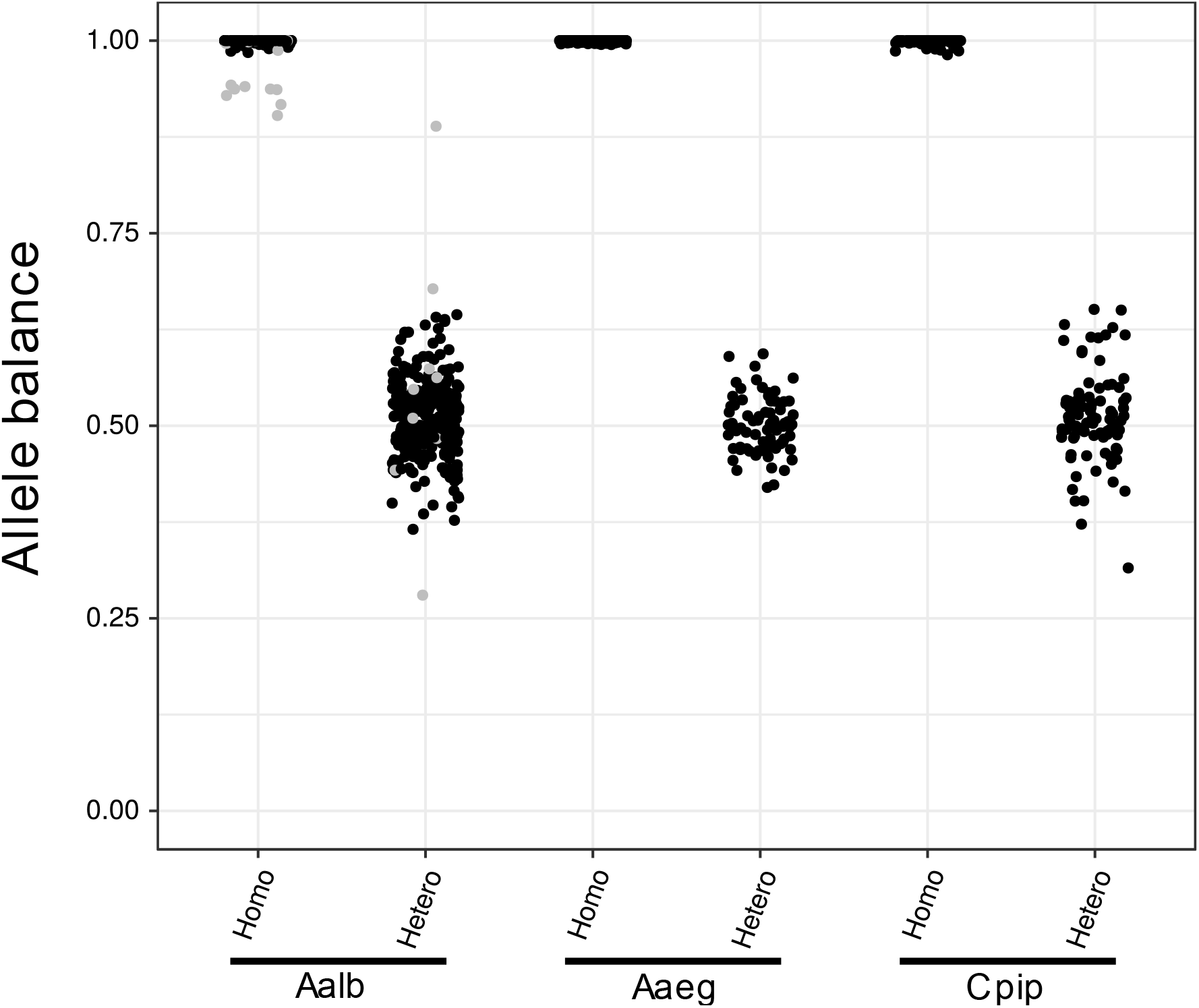
The allele balance in genotypes containing a variant. The distribution of allele balance in read depth for each allele at heterozygous or homozygous genotypes containing alternative allele(s). The balance was calculated as [read depth of the first allele in “GT” info] / [total depth] in the VCF format. Gray points in *A. albopictus* are at the genotype at GGT (Gly) trinucleotide repeats in exon32.

Aa substitutions identified at the end of the MoNaS pipeline are listed in Table 2. Of those, F1534C in Aalb-SP (Shinji Kasai et al., 2019), S989P and V1016G in Aaeg-SP (S Kasai et al., 2014), and L1014F in Cpip-JPP (Hardstone et al., 2007), all previously known to exist in those strains, were recalled correctly. In the Aaeg-Mex strain, V410L, V1016I, and F1534C variants, which are known as *kdr* (Brengues et al. 2003; Haddi et al. 2017; Kawada et al. 2009), were detected. Other aa substitutions detected—C749*Y (*in mosquito aa coordinates because there was no corresponding aa in *M. domestica*), A2023T and G2046E in *A. albopictus*, S723T in *A. aegypti*, and K109R, Y319F, T1632S, E1633D, E1856D, G2051A, and A2055V in *C. pipiens* complex—are not known for their effects on insecticide susceptibility.

**Table 2.** Detected amino-acid substitutions

| Species | Population | n | Amino-acid substitutions<br>(num. of homozygous, heterozygous individuals) |
| --- | --- | --- | --- |
| *A. albopictus* | Aalb-SP | 8 | F1534C**(8,0); A2023T(8,0) |
|  | Aalb-Viet | 2 | A2023T(0,1) |
|  | Aalb-Okayama | 2 | C749*Y(0,1); A2023T(0,1); G2046E(0,1) |
|  | Aalb-Toyama | 2 | A2023T(0,1) |
|  | Aalb-Ishigaki | 2 | A2023T(1,0) |
|  | Aalb-Yona | 8 | A2023T(0,2) |
| *A. aegypti* | Aaeg-Mex | 16 | V410L**(1,7); S723T(1,7); V1016I**(1,7); F1534C**(16,0) |
|  | Aaeg-SP | 8 | S989P**(8,0); V1016G**(8,0) |
| *C. quinquefasciatus* | Cpip-JNA | 2 | K109R(0,1); T1632S(0,1); E1633D(0,1);<br>G2051A(0,1); A2055V(0,1) |
| *C. quinquefasciatus* | Cpip-JPP | 2 | R261K(2,0); L1014F**(2,0) |
| *C. pipiens pallens* | Cpip-Ryo | 2 | Y319F(2,0); T1632S(0,2); E1633D(0,2) |
| *C. pipiens* form *molestus* | Cpip-JP_mix | 2 | Y319F(2,0); L1014F**(2,0); T1632S(2,0);<br>T1633D(2,0); E1856D(2,0) |

The amino-acid coordination is *M. domestica* except C749Y with asterisk (\*) since there was no corresponding amino-acid in *M. domestica* (genbank id: AAB47604). Double asterisks (\*\*) indicate known *kdr* substitutions conferring pyrethroid resistance. Variants seen on the Gly repeats near the C-terminal are omitted.

## Discussion

In this study, we evaluated potential of targeted enrichment sequencing of *VGSC* CDSs from the gDNA of mosquitos using hybridization capture probes. The result of the experiment is quite promising: most nucleotides of *VGSC* CDSs were covered at sufficient read depths even in samples with less than a 30 Mbp (0.1 million read-pairs) sequencing effort.

Even though the probe set was designed on the basis of the *A. albopictus* genome sequence only, it successfully enriched *VGSC* CDSs from the gDNA of two other Culicinae mosquito species, *A. aegypti* and *C. pipiens* complex, which are estimated to have diverged 71.4 and 179 million years ago, respectively, from *A. albopictus* (X.-G. Chen et al., 2015). Although the estimated entire efficiency of enrichment was lower by approximately 50% in *C. pipiens* samples than in *A. albopictus* samples, the on-target ratio was still comparable or rather higher in *C. pipiens* (Fig. 2). This result can be explained by the much smaller genome size of *C. pipiens* complex (579 Mb in the CpipJ2 assembly) as compared to *A. albopictus* (2.25 Gbp in the C6/36 assembly). Applicability of a single probe set to multi species (e.g., the same genus or family) is clearly advantageous because this obviates the need to prepare each custom probe sets specific for each single species and may enable capture even in species lacking prior genome information. Nonetheless, the evolutionary distance will limit the range of species that one probe set can be applied. In this study, the mapping results on each exon indicated that capture efficiency decreased in some exons (Fig. 3). The empirical observation suggests that less than 87.5% in identity or less than 60 bp in size for the homology track of targets could decrease the efficiency of capture significantly (Fig. 5). Especially, our probe set failed to capture two optionally used exons, 2 and 16.5, in *A. aegypti* and *C. pipiens* complex, which are among the smallest exons targeted (Fig. 3). It is assumed that those tiny exons alone do not provide enough thermostability for probe–target DNA duplex during the capture. Because the probes for those small exons contain flanking intronic sequences of the *A. albopictus* genome, those flanking sequences may have provided enough homology region to capture sequences from this species. Although it is straightforward to optimize our probe set further at least for the two other species of mosquito simply by adding species specific probes for those exons and flanking intronic regions, small exons in general will be a major challenge when a probe set is aimed to be used for a group of species rather than specific targets because the homology in an intronic region will decay more rapidly than that in an exonic region during speciation. We also missed another exon corresponding to “exon 12” in *Anopheles gambiae* described by Davies et al. (2007) during probe design (see Materials and Methods). Such tiny and rarely used exons may be difficult to annotate without high-quality high-throughput RNA sequencing data. Nevertheless, in mosquitoes, all such tiny exons are actually situated on the N-terminal intracellular loop or the intracellular loop between domains I and domain II in VGSC, where no resistance-associated mutation has been found so far (Dong et al., 2014). Therefore, it is not clear whether ignoring those small exons of *VGSC* from analysis does pose a serious problem for insecticide resistance research.

**Fig. 5.**
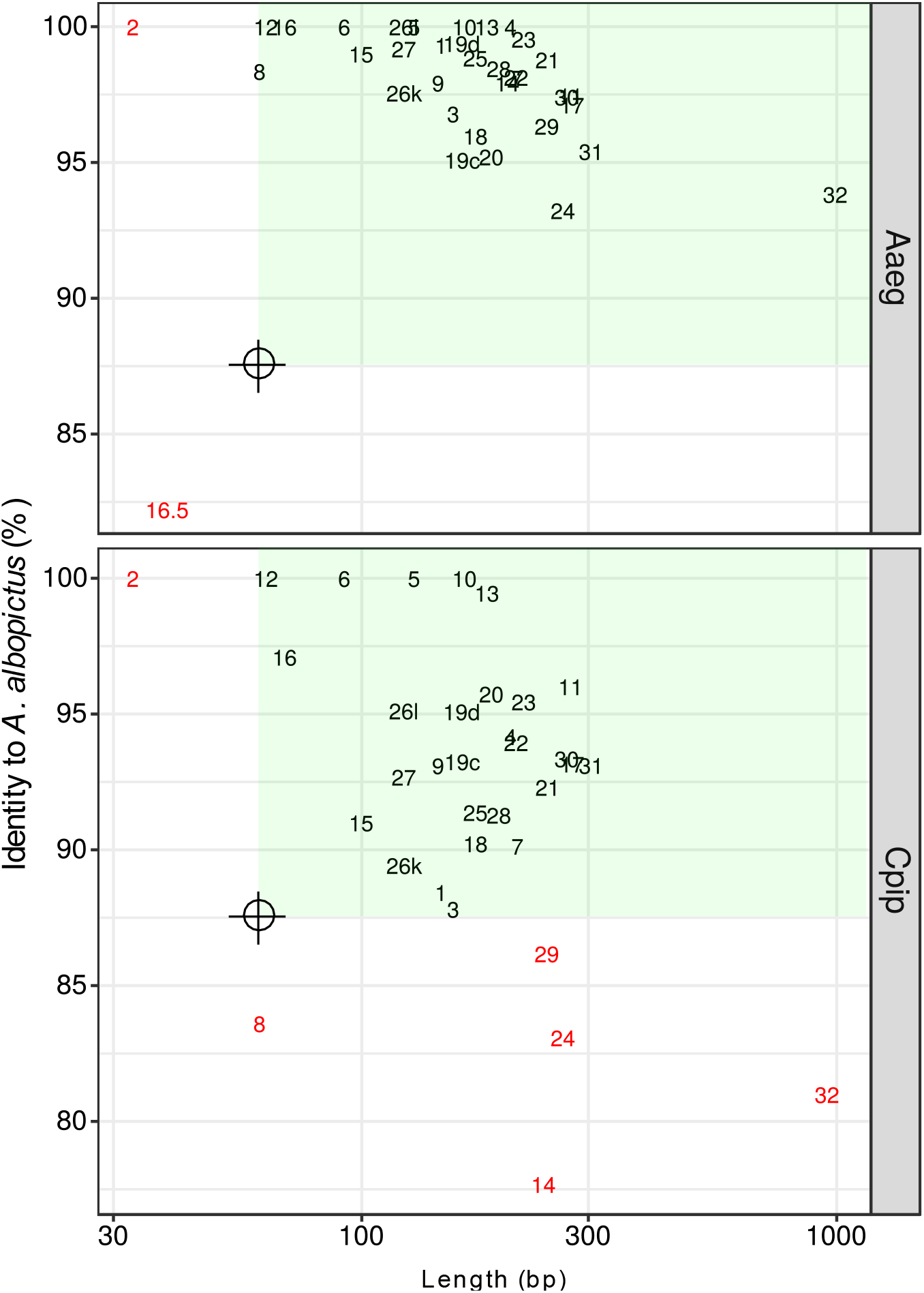
Length and conservation of the *VGSC* exons (CDSs) targeted. Length (on a logarithmic scale) and percentage identity to *A. albopictus* of each exon in *A. aegypti* and *C. pipiens* complex. Red exon names are those with low-coverage nucleotides in Fig. 3. The green area represents >60 bp length and >87.5% identity.

The process of genotyping *VGSC* carried out in this study was automated in MoNaS. This program sequentially runs tools conducting mapping of NGS reads to a reference, sorting, removal of PCR duplicates, indexing for BAM files, variant calling, variant annotation, and finally integration of these results across multiple samples into a single table with conversion of the aa coordinates to those corresponding to the *M. domestica* VGSC protein. The automation in MoNaS allows researchers to process raw NGS reads of many samples via a simple command line operation without expert knowledge of the bioinformatics field. MoNaS can be run locally with appropriate genome reference data. Also, a web-service of MoNaS implemented with JBrowse alignment viewer (Buels et al., 2016) is provided by NIID Pathogen Genomics Center’s severer (https://gph.niid.go.jp/monas) (Fig. 1B).

## Supporting information

S3 Appendix.zip

## Acknowledgements

This research was partially supported by the Research Program on Emerging and Re-emerging Infectious Diseases (JP19fk0108067) from Japan Agency for Medical Research and Development (AMED) and by Japan Initiative for Global Research Network on Infectious Diseases (J-GRID) from the Ministry of Education, Culture, Sports, Science and Technology of Japan and AMED.

## Authors’ contributions

KI and SK designed the study. KI and OK designed the capture probes. KI and TT refined the annotation of the *VGSC* gene. KI conducted all the experiments. KI, TSe, KY, and MK contributed to the bioinformatic analysis and development of MoNaS. SK, YM, MS ans TSa contributed maintaining the laboratory colony or collected specimens of mosquitoes in the field. KI drafted this manuscript. All the coauthors critically revised the manuscript and approved it for publication.

## Data accessibility

Raw NGS reads obtained in this study were deposited to DDBJ Sequence Read Archive (BioProjectID: PRJDB7889, see Table 1). Annotation information for *VGSC* and new reference sequence of *VGSC* gene in *C. pipens* complex used in this study (supercont3.182_3) are provided in S3 Appendix.zip file. Web service and source codes of MoNaS are hosted in https://gph.niid.go.jp/monas and https://github.com/ItokawaK/MoNaS, respectively.

